# Gene regulation by synergism between mRNA decay and gene transcription

**DOI:** 10.1101/2022.04.19.488766

**Authors:** José García-Martínez, Abhyudai Singh, Daniel Medina, Sebastián Chávez, José E. Pérez-Ortín

## Abstract

It has become increasingly clear in the last few years that gene expression in eukaryotes is not a linear process starting from transcription in the nucleus to the cytoplasm, but a circular one where the mRNA level is controlled by crosstalk between nuclear transcription and cytoplasmic decay pathways. One of the possible purposes of this crosstalk is to keep the mRNA level approximately constant. This is called mRNA buffering and happens when transcription and mRNA degradation act at compensatory rates. However, if transcription and mRNA degradation act synergistically, enhanced gene expression regulation would occur. In this work we mathematically modeled the effects of RNA binding proteins (RBP) when they have positive or negative effects on mRNA synthesis and decay rates. We found that they can buffer or enhance gene expression responses depending on their respective effects on transcription and mRNA stability. Then we analyzed new and previously published genomic datasets obtained for several yeast mutants related to either transcription or mRNA decay, but they are not known to possess activity in the other process. We show that some of them, which were presumed only transcription factors (Sfp1) or only decay factors (Puf3, Upf2/3), may represent examples of RBPs that make specific synergistic crosstalk to enhance the control of the mRNA levels of their target genes by combining antagonistic effects on transcriptional and mRNA stability.

## Introduction

It has been postulated that crosstalk between transcription and degradation machineries acts to control mRNA concentration ([mRNA], Sun et al., 2012; 2013; García-Martínez et al., 2021a). This crosstalk has two branches: transcription to degradation (direct: from the nucleus to the cytoplasm) and degradation to transcription (reverse: from the cytoplasm to the nucleus; see Figure 1), which converts gene expression into a circular system (Haimovich et al., 2013; Medina et al. 2014). The mechanisms and purposes of both directions can differ. For the direct direction, the mechanism has been demonstrated as the co-transcriptional imprinting (Choder 2011) of mRNAs with RNA binding proteins (RBPs). mRNA imprinting has been shown in the yeast *Saccharomyces cerevisiae* for: RNA polymerase II subunits Rpb4/7 (Goler-Baron et al., 2008; Dori-Bachash et al., 2011, Shalem et al., 2011; Richard et al, 2020); basal transcription machinery factor Taf7 (Cheng et al, 2021); the Ccr4-NOT complex (Villanyi et al, 2014; Collart & Panasenko 2017); specific decay factors Cth2 (Vergara et al., 2011) and Puf3 (Garrido-Godino et al, 2021). Imprinting RBPs should return back to the nucleus later. This return to the nucleus can be used as a reverse direction mechanism (Figure 1, see below). Note that most of these RBPs have been originally described as part of either transcriptional machinery (Rpb4/7, Taf7) or degradation pathways (Cth2, Puf3), or both (Xrn1, Ccr4-Not). mRNA imprinting can have compensatory effects by increasing or decreasing synthesis rates (SRs) and decay rates (DRs) and, at the same time, causing mRNA buffering. Buffering can be global by acting on the whole mRNA concentration ([mRNA]_t_) or being restricted to a subset of genes depending on the fraction of the transcriptome bound by RBPs.

**Figure 1.**
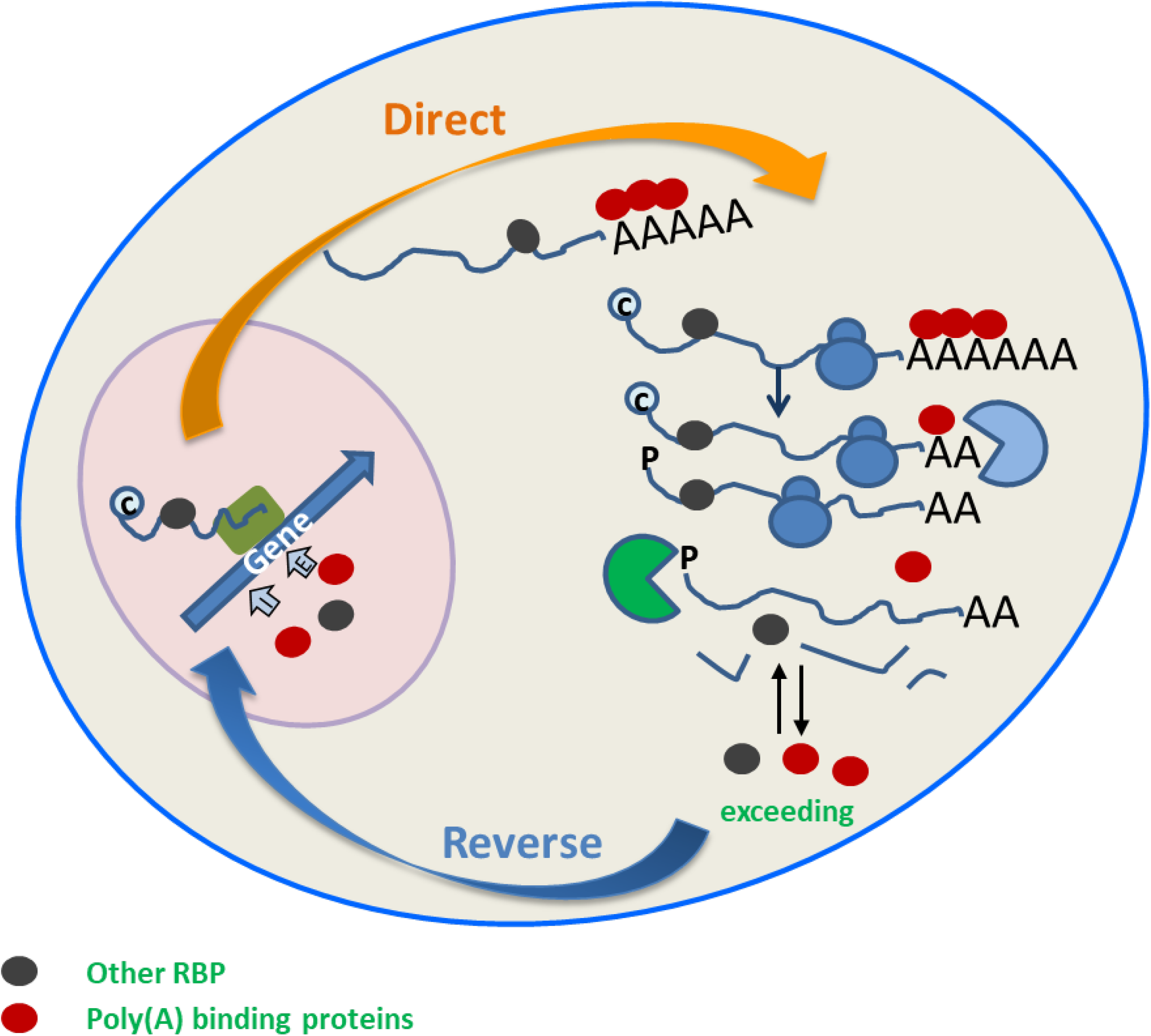
Transcription-degradation crosstalk pathways. Crosstalk has two branches: direct from synthesis (transcriptional) machinery for the mRNAs in the nucleus to the decay machineries in the cytoplasm; reverse from the cytoplasm to the nucleus. RNA binding proteins (RBP, either polyA-binding or others) can co-transcriptionally imprint mRNAs and be transported with them to the cytoplasm where they affect the mRNA life, especially mRNA stability. Once mRNA is degraded, they can be imported back to the nucleus where they affect transcription. See the main text for further explanations.

Global [mRNA]_t_ buffering has been demonstrated to occur when subunits of transcriptional or decay machineries are depleted. In the yeast mutants that lack some decay subunits, increased mRNA stabilities (decreased DRs) are balanced with lower global mRNA synthesis rates, which maintain the steady-state levels of most mRNAs stable (Sun et al., 2013; Haimovich et al., 2013). Similarly in other yeast mutants that lack subunits of transcription complexes, decreased SRs are balanced by increasing mRNA stabilities (Timmers & Tora, 2018). The molecular mechanisms underlying this balancing process are not well understood. As previously explained, direct crosstalk may be based on mRNA imprinting with the RBPs that modulate mRNA stability later in the cytoplasm (Choder 2011). Reverse crosstalk needs a mechanism to not only send information from the cytoplasm to the nucleus (see below), but to also convert this information into a transcriptional effect. One possible mechanism for this is the regulation of transcription elongation by RBPs. Slobodin et al, (2020) found that regulation at this level impacts the length of poly(A) tails and the expression of decay machinery (Ccr4-NOT) as a mechanism for [mRNA]_t_ buffering. The effect of the Ccr4-NOT complex as a stimulator of transcription elongation has also been demonstrated in yeast (Collart and Panasenko, 2017; Reese, 2013). We also found that the effect of Xrn1 on synthesis rates in yeast is executed, at least in part, by favoring the elongation rates of RNA polymerases (Haimovich et al., 2013) in parallel to Ccr4-NOT (Begley et al, 2019). This occurs by preventing backtracking (García-Martínez et al., 2021b), especially near the poly(A) site (Begley et al. 2021; Fischer et al., 2020).

The reverse crosstalk direction also needs a mechanism to send information from the cytoplasm to the nucleus, as previously stated. This can be done by the nuclear import of the cytoplasmic proteins coupled with the direct crosstalk direction to bring them back to the cytoplasm. This idea has been used in the models proposed by B. Glausinger and T. Jensen (Gilbertson et al., 2018; Schmid and Jensen 2018). These models propose poly(A)-binding proteins as sensors of the level of cytoplasmic poly(A) mRNAs. Interestingly, the reverse crosstalk in the study of Gilbertson et al. (2018) does not act by activating transcription as a buffering system to compensate for an increase in mRNA decay but, on the contrary, to enhance the drop in most mRNA levels. This example shows that transcription/degradation crosstalk can be not only compensatory for mRNA buffering, but also synergistic to reinforce a primary regulatory stimulus (Hartenian & Glaussinger, 2019).

The consequence of crosstalk can also differ when acting on a limited number of mRNA targets. The crosstalk direction from the nucleus to the cytoplasm may have the effect of determining the future cytoplasmic life of some mRNAs according to which environmental circumstances they were synthesized in. For instance, the mRNAs transcribed during stress responses can be imprinted to become more stable to more quickly increase cytoplasmic levels (Slobodin et al., 2020) or, on the contrary, with less stability to provoke a sharper response peak that reduces the cost of the response (Pérez-Ortín et al, 2007; Shalem et al., 2011). Sending information from the cytoplasm to the nucleus for a limited set of mRNAs has been shown in the phenomenon called genetic compensation in higher eukaryotes. During this process, the mRNAs bearing premature stop codons (PTCs) are degraded by the usual non sense-mediated decay (NMD) route for PTCs mRNAs, but with a higher synthesis rate of other sequence-related paralogous genes that partially compensates their mRNA levels (Ma et al 2019; El-Brolosy et al. 2019). This can partly compensate the phenotypic defect caused by the primary mutation. This mechanism is not, however, universal because it does not apparently exist in *S. cerevisiae* (García-Martínez et al., 2021a).

We reasoned that in order to investigate the existence of both crosstalk directions in a set of mRNAs, it is necessary to study the factors that have been described to control a group of genes at either the transcriptional or the mRNA stability level. In this study, we first mathematically modeled the different possibilities of transcription-mRNA decay crosstalk by the RBPs bound co-transcriptionally in the nucleus. We found that mRNA imprinting by RBPs is a very efficient way to buffer [mRNA] when RBPs have parallel (stimulating or inhibitory) effects on synthesis and degradation rates. We also found that upon having a synergistic effect, the activity of RBPs at the transcription and post-transcriptional levels speeds up and enhances the response, which suggests that they could be useful as a regulatory mechanism for eukaryotic cells. Then to check the various theoretical possibilities predicted by modeling, we used some new and previously published genomic datasets of several yeast mutant strains of the model eukaryote *S. cerevisiae*. These strains are depleted of proteins, previously known to be related to either transcription or mRNA decay, but not known to possess activity in the other process. Here we show that some RBPs, which are probably exported and bound to mRNAs from the nucleus, seem to synergistically enhance the synthesis and decay rates of their target mRNAs. This occurs when the protein is both a presumed transcription factor (Sfp1) or a presumed decay factor (Puf3, Upf2/3), and suggests that these RBPs make specific crosstalk to control the mRNA levels of their targets by combining transcriptional and mRNA stability effects.

## Results

### Mathematical modeling of mRNA levels based on transcription/decay crosstalk pathways

We decided to model the behavior of the RBPs with either mRNA stabilizing or destabilizing activity, and to also show the ability to influence transcription either positively (activator) or negatively (repressor). For this purpose, we used an ordinary differential equation-based mathematical model to capture the biomolecular crosstalk circuit dynamics, which allowed us to predict the mRNA level outcome according to the synthesis rate (Figure 2A). We assumed that RBP binds mRNA co-transcriptionally, and can affect the initiation and/or elongation of RNA pol II based on our previous results with the Xrn1 and Ccr4 factors (Begley et al, 2019). We also assumed that the bound RBP factor travels with mRNA to the cytoplasm and remains bound to it until mRNA is degraded (Chattopadhyay et al., 2021).

**Figure 2.**
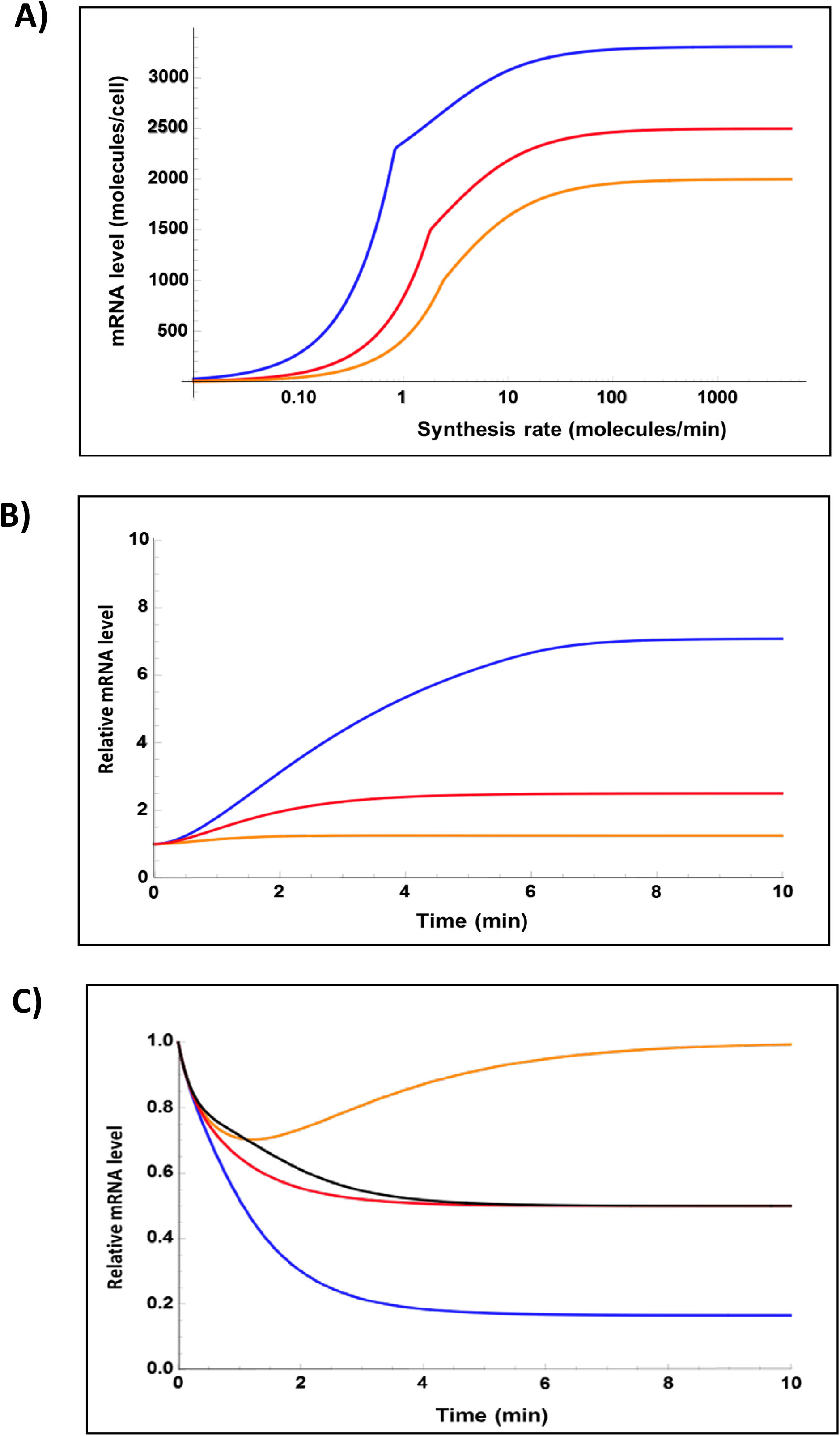
Mathematical modeling of the transcriptional responses conducted by crosstalk factors. A) mRNA response as a function of the synthesis rates for the case of a transcriptional activator with stabilizing (blue line) or destabilizing (orange line) activity compared to the case in which the factor has no mRNA binding capacity (red line). B,C) Modeling different response curves during transcription activation (B) or repression (C) for the cases of the RBPs that possess parallel or antagonist activities at the SRs and DRs. Transcriptional activation means turning on a transcriptional activator or turning off a transcriptional repressor. Repression means the opposite in both cases. Blue curves are for the cases with a transcription activation factor that possesses stabilizing activity. A transcriptional repressor with destabilizing activity would show similar plots. Orange curves are for the RBP transcription factors with opposite effects on mRNA stability and provoke buffering. There is no crosstalk for a TF with no mRNA binding properties (red curves) or for a destabilizing factor with no transcriptional activity (black curve). The time scale shown is the case of an mRNA with a HL of 1 min as a reference.

We modeled the cases of the factors that activate transcription and enhance mRNA degradation (i.e., Rpb4/7; Xrn1, Ccr4-NOT, etc.) compared to simple activators, which do not influence mRNA stability (see M&M for a detailed description). Figure 2A depicts not only how such an activator (orange line) is less efficient in increasing mRNA levels at any synthesis rate, but how the maximum mRNA level possible is lower than that obtained with an activator without mRNA binding activity (red line). If the activator possesses mRNA stabilizing activity (blue line), it serves to obtain higher mRNA levels for any synthesis rate. As far as we know, such transcriptional activators have not yet been described. The case of transcriptional repressors possessing mRNA binding activity and effects on stability is similar to that of activators, but it considers that gene activating occurs when the repressor is turned off and when gene repression is turned on. The obtained lines are similar to those shown in Figure 2A (not shown).

Having seen the enhancement caused by RBPs, which act synergistically on mRNA levels upon transcription and mRNA stability, we wondered what would happen during the kinetics of the activation or repression of a gene (or a group of co-regulated genes) regulated by this kind of RBPs. In Figure 2B-C, we compare these results (blue lines) to the cases with no mRNA binding activity (red and black lines), and also to the cases of antagonist effects (orange lines). During activation, the RBPs that play dual activities have strong effects on both the kinetics and final levels of mRNAs (Figure 2B). If the transcription activator is also an mRNA stabilizing factor, mRNA levels more quickly increase and reach higher levels than when it acts only at the transcription level (compare the blue and the red lines in Figure 2B). Similar results are obtained if a factor that acts as a transcriptional repressor and destabilizes mRNA is switched off. In contrast, we obtain buffering behavior (orange line) when the transcription activator induces mRNA degradation.

We also modeled the action of the same series of factors during the repression kinetics (Figure 2C). When a repressor with mRNA destabilizing activity is switched on (or an activator is switched off), repression is faster and stronger if the repressor has synergistic activities on mRNA stability (blue line) than when it acts only at the transcription level (red line). Similar results are obtained when an RBP possesses only mRNA destabilizing activity and not a transcriptional repressor (black line). Buffering is obtained (orange line) when the activator possesses mRNA destabilizing activity after being switched off.

All the possible kinds of such RBPs can be called crosstalk factors (CFs) because they serve to send information from the nucleus to the cytoplasm by means of mRNA imprinting. Depending on their activities on chromatin and in the cytoplasm, they can serve as buffering factors (BFs) or, alternatively, to speed up transcriptional responses as regulatory factors (RFs). In the case of [mRNA] buffering, we previously showed that it is mostly global in the yeast *S. cerevisiae* by controlling total [mRNA], and not the concentration of particular sets of mRNA. However, with RFs, it seems logical that they can be devoted to specific groups of genes. Therefore, we decided to look for them in the published datasets of synthesis rates and mRNA stabilities of *S. cerevisiae*.

### Looking for examples of synergistic crosstalk in post-transcriptional regulons

In order to check the existence of the predicted crosstalk modes based on RBPs, we ran a meta-analysis of a previously published set of yeast deletion mutants, where each one lacked a factor related to a pathway involved in mRNA decay (Sun et al., 2013). In that study, the existence of global [mRNA]_t_ buffering was established for almost all the analyzed mutants (Sun et al, 2013).

This dataset included some examples of RBPs with specific sets of targets, e.g. the Puf and Upf proteins that have been related to either mitochondria (Puf3), ribosome biogenesis (Puf4) or cell periphery (Puf1/2) (Gerber et al., 2006), or the preferred mRNAs of the NMD pathway (Upf2/3) (Brogna et al, 2016). We analyzed the relative behavior of the targets in mRNA, half-lives (HLs) and SRs. As expected, the relative HLs of their targets clearly increased for mutants *puf3, upf2* and *upf3*, along with relative increases in SRs, which resulted in a synergistic effect upon which the mRNA levels of their targets increased by lowering decay and raising transcription (Figure 3A-C). *puf4*, however, showed a significant effect on the HLs of its targets, but not at the transcription level (Figure 3D). This indicates that it is not a crosstalk factor. Finally, *puf1* and *puf2* show no clear effects on the mRNA stability of their targets in spite of their presumed activity (Figure 3E-F), while *puf1* has a very weak synergistic effect on the SR.

**Figure 3.**
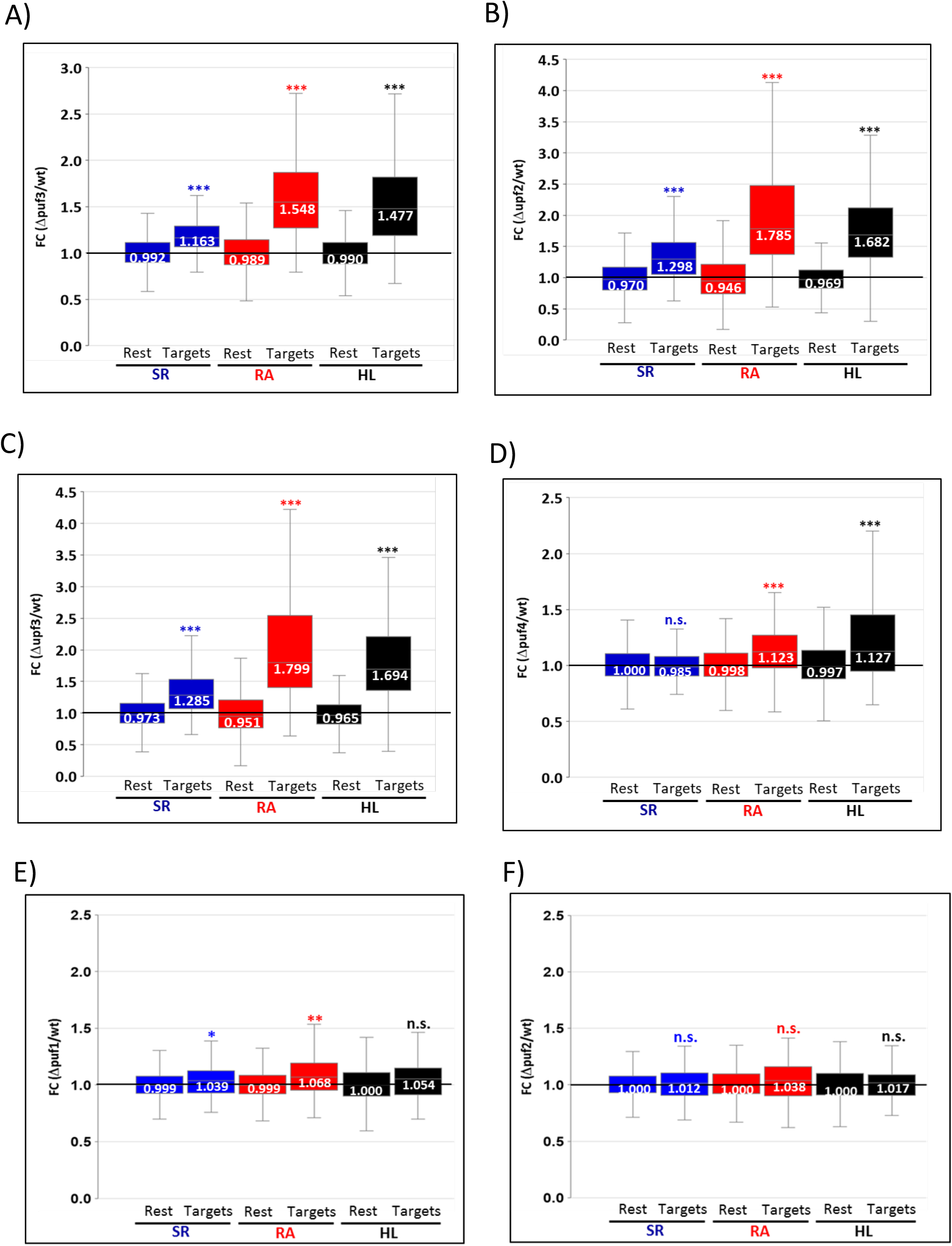
Transcriptomic study of several yeast strain mutants in mRNA decay factors. Fold changes (FC) for the synthesis rates (SR), mRNA half-lives (HL) and mRNA levels (RA) in the mutants *vs*. the wild type strains are represented in the box-plots. We compared the median of the FC of the targets for Puf1 and Puf2 (70 genes), Puf3 (210 genes), Puf4 (145 genes), Upf2 and Upf3 (479 genes) to the rest of the transcriptome. The results show the synergistic effect of the Puf3 (A), Upf2 (B) and UPF3 (C) factors. However, Puf4 (D) only affects the HLs of its targets, whereas Puf1 (E) and Puf2 (F) have very slight effects. The numbers in boxes show the median value of the FC. The statistical significance for the pair-wise comparisons (targets vs. the rest) was estimated by the Kolmogorov-Smirnov test of the differences between distributions (*** is p-value <10^−5^; ** p-value < 0.005; * p-value <0.05; n.s. not significant).

These results reveal that some RBPs, which have initially been presumed to affect only the mRNA stability of their targets (e.g. Puf3, Upf2, Upf3), also act in transcriptional repression for those genes demonstrating the existence of specific crosstalk, which seems to synergistically act by summing up effects on SRs and DRs as a way to increase the levels of a group of RBP target mRNAs.

### Crosstalk mRNA transcription/decay in transcriptional regulons: studying the *sfp1* mutant

We decided to extend our analysis to transcriptional activators. To this end, we used the case of the yeast Sfp1 factor. Previously published works reveal that steady-state mRNA measurements in *sfp1Δ* strains show the up- or down-regulation of a relatively large numbers of genes (Jorgensen et al. 2004; Cipollina et al. 2005), which makes it a good candidate to explore the existence of specific crosstalk. Sfp1 is a transcription factor that primarily binds the promoter of ribosomal proteins (RPs) and affects ribosome biogenesis (Reja et al., 2015).

We performed a GRO analysis to elucidate the effect of Sfp1 depletions on SR, RA and HL parameters. The comparison of Sfp1 targets against the effect on the rest of the transcriptome (Figure 4) showed in *sfp1Δ* that, as expected, there was a strong effect on the RNA pol II SR of their RP targets as regards the global average. There was, however, a stronger effect on their mRNA levels. This was caused by destabilization, which occurs in RP mRNAs (Figure 4). These results indicate that Sfp1 activates all its RP targets at the transcription level, but it also has a synergistic effect by increasing their mRNAs’ stability, which contributes to the final mRNA levels.

**Figure 4.**
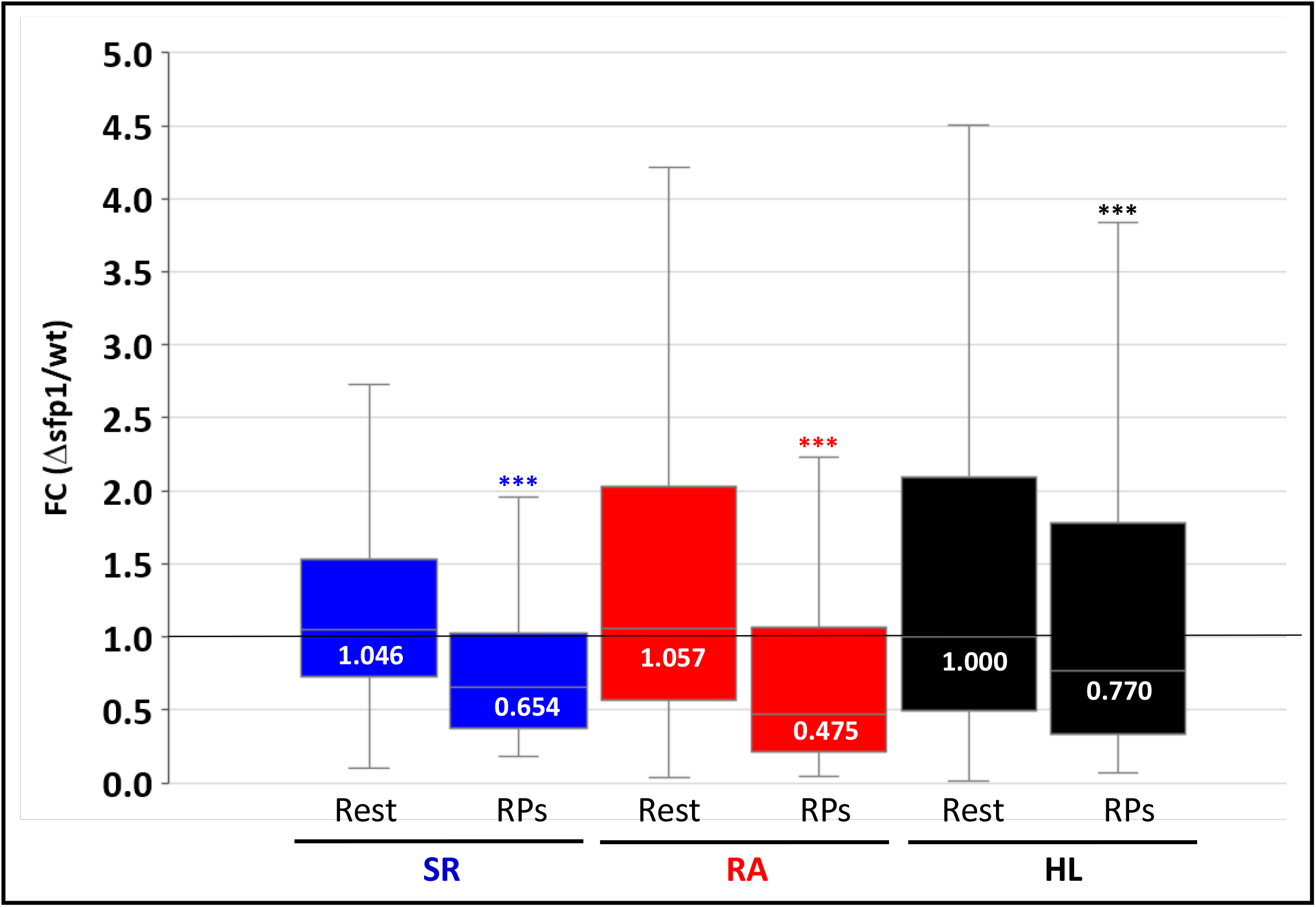
Transcriptomic analysis of the *sfp1* mutant. We performed a GRO analysis of the *sfp1* mutant compared to its wild-type strain (BY4741). Then we compared the fold changes of the group of 106 RP Sfp1 targets to the rest of the transcriptome as explained in Figure 3. The experiment was run in triplicate. The numbers in boxes show the median value of the ratio (FC). The statistical significance for the pair-wise comparisons (targets vs. the rest) was estimated by the Kolmogorov-Smirnov test of the differences between distributions. *** is p-value <10^−5^.

## Discussion

mRNA transcription/degradation crosstalk is a process that, in spite of having been quite recently discovered (see Pérez-Ortín et al, 2013 for a review), is attracting plenty of attention because it probably influences all aspects of eukaryotic gene expression. The features and mechanisms of this crosstalk are, however, much less known. In this study we wondered whether mRNA buffering is the only outcome of these crosstalk mechanisms or if they can also contribute to regulate changes in gene expression. We modeled the effect on the mRNA levels caused by the regulators that act in the same or opposite direction (activation or repression) in both transcription and mRNA stability. We predicted that when acting in the same direction, such complex regulators would enhance changes in mRNA levels by favoring faster and stronger regulatory responses. By doing so, they would contrast with the crosstalk factors involved in buffering, which display opposite effects in transcription and mRNA decay.

The expected result for a regulon would be the SR of the genes activated by the depleted transcription factor to lower and, because of this, so would their mRNA levels. The expected result for a post-transcriptional regulon is that the mRNAs bound by an RBP, which induces their degradation, would increase their HLs and, consequently, their mRNA levels. If specific crosstalk exists for these mRNAs, the observed changes in their levels would differ by either raising or lowering from global mRNA changes. The relative change in the average mRNA levels of the genes that behave with the regulon, compared to the change in the global mRNA levels, would inform us if there was i) specific compensatory crosstalk to buffer regulon mRNA levels, ii) synergistic crosstalk that exacerbates the effect on those mRNAs or iii) no specific crosstalk.

We investigated seven previously described post-transcriptional regulons driven by RBPs (Upf1/2/3 and Puf1/2/3/4). In those mutants, a previous study showed that global buffering was not affected (Sun et al., 2013). In four of them, we confirmed that the HLs of their target mRNAs had increased (*puf3, puf4, upf2, upf3*). When we analyzed the transcriptional effect on their target genes, we found no effect on *puf4* (Figure 3D), which indicates that it is a simple regulator that operates only at the post-transcriptional level. This case is similar to the previously studied one of *upf1* (see García-Martínez et al. 2021a). Nevertheless, and quite unexpectedly, we found synergistic behavior for three other studied cases: *puf3, upf2* and *upf3*. In all three cases, we concluded that these RBPs not only stimulate the decay of a selected group of mRNAs, but also play a negative role in their transcription. Therefore, the relative increase in the mRNA levels observed in their target mRNAs is due to not only the disappearance of a specific decay factor, but also to lack of specific transcriptional repression. We do not know what mechanism of transcription repression these RBPs follow, and if it is direct or indirect, but we recently found that Puf3 is also a nuclear protein that binds chromatin and imprints its mRNA targets (Garrido-Godino et al, 2020). For the Upf2/3 proteins, nuclear location has been demonstrated in human cells, where hUpf3 shuttles between the cytoplasm and the nucleus, hUpf2 is perinuclear and hUpf1 is mainly cytoplasmic (Lykke-Andersen et al, 2000). The results are less clear in yeast, but Upf3 has an NLS (nuclear location signal; see SGD: https://www.yeastgenome.org/). The results herein obtained for *upf2/3* contrast with those obtained for *upf1* (García-Martínez et al. 2021a). This result is, at first glance, surprising given the presumed function of all three proteins in the NMD pathway that forms a protein complex (Gupta & Li, 2018; He & Jacobson 2015). The functions of these three proteins can, however, be multiple and differ from one another. Indeed, it has been demonstrated that hUpf1 can be imported to the nucleus where it plays a role in nonsense-mediated altered splicing (NAS), a pathway that probably does not exist in yeast, which indicates that putative nuclear functions associated with Upf1 emerged recently during evolution and might be a unique feature of the mammalian Upf11 protein (Azzalin 2012). Thus if yeast Upf1 is only involved in cytoplasmic NMD, but Upf2/3 perform additional nuclear functions, only the last two Upf proteins may affect transcription and imprint mRNAs similarly to Puf3. In fact, it has been demonstrated that Upf machinery interacts at the nucleus with epigenetic machinery in higher eukaryotes, which provides a link between the cytoplasm and the nucleus for this response (El-Brolosy et al. 2019; Ma et al. 2019).

For Puf3, whose targets are mRNAs that encode mitochondrial proteins (Quenault et al, 2011), synergistic action on synthesis and degradation rates could contribute to a faster down-regulation of the mRNA levels of targets (as modeled in Figure 2C: blue vs. black lines) when cells undergo fermentation conditions, and act repressively upon both mRNA synthesis and decay.

We also wished to analyze an example of a well-known activator that drives a transcriptional regulon. We analyzed by Genomic Run-On the *sfp1* mutant strain following the same principle previously described for post-transcriptional regulons. The results we obtained with the Sfp1 regulon indicate the existence of a synergistic effect on the RP mRNAs mediated by their co-transcriptional imprinting. The influence of a transcription factor (TF) on the mRNA HL that it activates has been demonstrated in yeast (Bregman et al, 2011; Trcek et al, 2011; Zander et al, 2016). In such cases, specific *cis* elements of the promoter, which are bound by specific TFs, direct the imprinting of transcribed mRNAs that are either more quickly exported (Hsf1-dependent heat-shock mRNAs, Zander et al, 2016) or accelerate decay (Rap1-dependent mRNAs, Bregman et al, 2011; and cell cycle-regulated mRNAs during mitosis, Trcek et al, 2011). Imprinting can be done by the same proteins that bind the promoter. One such case is Dbf2 for cell cycle mRNAs (Trcek et al., 2011) or, alternatively, the TF can direct mRNA imprinting by a more general factor like the Mex67 export protein in heat-shock mRNAs (Zander et al, 2016).

Based on the results of this study and those in a previous paper (García-Martínez et al, 2021a), we propose a model based on the imprinting mechanism by RBPs to explain the direct crosstalk direction. Imprinting might vary from very specific to global depending on the nature of the RBPs, which can bind a variable number of mRNAs. RBPs affect the stability of their targets in the cytoplasm and should then return to the nucleus (Figure 1). The nuclear import of free RBPs would be the mechanism for the reverse crosstalk direction (Chattopadhyay et al., 2021). We demonstrate here that this would be an efficient way to buffer [mRNA]_t_ if RBPs possess transcriptional activation activity, but can also be a mechanism for the synergistic effect if RBPs negatively affect transcription and mRNA decay, such as Puf3 and Upf2/3. Possible models for the mechanism of reverse global crosstalk have been published. They are based on the nucleo-cytoplasmic shuttling of a kind of wide-binding spectrum RBPs: polyA binding proteins. Glaussinger’s group (Gilbertson et al, 2018), proposed an anti-buffering mechanism mediated by the cytoplasmic poly(A) binding protein (PABPC), whereas Jensen’s group proposed a buffering mechanism based on the Nab2 function in nuclear mRNA decay (Schmid & Jensen, 2018). RBPs are the best candidates to shuttle information forward and backward from the nucleus to the cytoplasm. They can sense the [mRNA]_t_ in the cytoplasm and be imported to the nucleus whenever they are free, where they can have positive or negative effects on transcription (Hartenian & Glaussinger, 2019). The existence of proteins that are simultaneously gene transcription factors, bind DNA, play roles in mRNA metabolism and bind mRNA is becoming a frequent finding (Conrad et al, 2016; Xiao R et al, 2019).

Synergistic crosstalk promotes regulatory changes whereas compensatory crosstalk ensures mRNA buffering (Figure 5). These two mechanisms operate in opposite directions. As buffering operates globally (García-Martínez et al, 2021a), it should act as a general obstacle for gene regulation, particularly when a large number of genes have to coordinately change their expression significantly. Under these conditions, slow regulation might be insufficient to counteract robust global buffering mechanisms. According to our mathematical modeling, synergistic changes by simultaneous actions upon mRNA synthesis and decay would involve a kinetic advantage to overturn buffering via a mechanism of signal amplification. From this point of view, crosstalk mechanisms would contribute to global [mRNA] homeostasis by diminishing stochastic noise (compensatory action, buffering) and, at the same time, favoring fast regulations in specific regulons in response to meaningful regulatory signals (synergistic-amplified regulation) (Figure 5). Hence it is interesting to note that here we show how Sfp1 synergistically regulates RPs genes, which are highly active under fermentative growth conditions, whereas Puf3 regulates mitochondria-related genes, which are expressed under respiratory conditions. In both cases, the activation or repression of the regulon should compensate for the general buffering mechanism that occurs when the carbon metabolism changes (García-Martínez et al., 2004). Interestingly, we observed how these two regulons feature among the few that escape from global [mRNA] buffering during the growth rate variation caused by transcription-mRNA decay crosstalk (García-Martínez et al., 2016).

**Figure 5.**
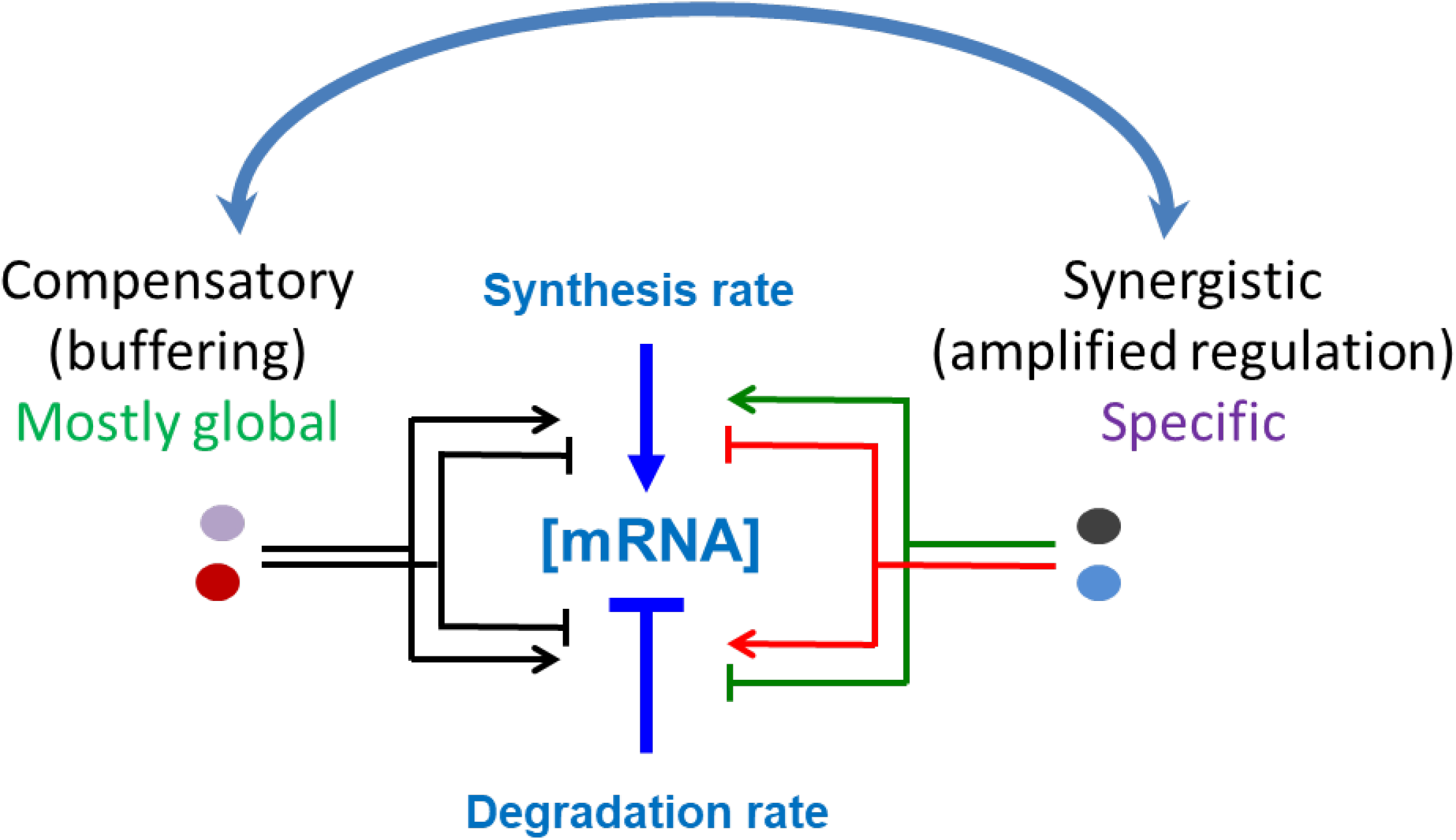
Summary of compensatory or synergistic effects in the transcription-mRNA decay crosstalk mediated by RNA binding proteins. Crosstalk can affect the respective SRs or DRs by increasing them synergistically, which sums up the effect of both changing (raising or lowering) the mRNA level [mRNA] (right). Alternatively, crosstalk can act by compensating the effects of SR and DR changes in a way that buffers [mRNA] (left). mRNA buffering normally affects global [mRNA] (Sun et al., 2013; García-Martínez et al., 2021a), whereas synergistic crosstalk acts on specific subsets or co-regulated genes by provoking an amplification of their regulation (see Figure 2). The existence of global [mRNA] buffering and specific synergistic regulation of a subset of mRNAs should involve a mutual influence (upper blue arrow).

## Materials and Methods

### Yeast strains and culture media

*sfp1Δ* yeast (the Y05352 strain from the Euroscarf collection: BY4741 *sfp1Δ*::KanMX4) and wild-type (BY4741: *Mat a, leu2Δ, his3Δ, met15Δ, ura3Δ)* cells were grown in YPD medium (1% yeast extract, 2% peptone, 2% glucose) at 30ºC. Pre-cultures were grown overnight. The next day, pre-cultures were diluted to OD_600_ = 0.05 to be grown in 250 mL flasks with agitation at 190 rpm until an OD_600_ of ∼0.5 was reached. Cells were recovered by centrifugation and flash-frozen in liquid nitrogen.

### Genomic methods

Genomic run-on (GRO) in the *sfp1Δ* mutant was performed as described in (García-Martinez et al. 2004), and as modified in (García-Molinero et al., 2018). Briefly, GRO detects, by macroarray hybridization, genome-wide active elongating RNA pol II, whose density per gene is taken as a measurement of its synthesis rate (SR). At the same time, the protocol allows the mRNA amounts (RAs) for all the genes to be measured. mRNA half-lives (HLs) are calculated as RAs/SRs by assuming steady-state conditions for the transcriptome.

### Modeling

We use the following ordinary differential equation systems to model the crosstalk circuit:

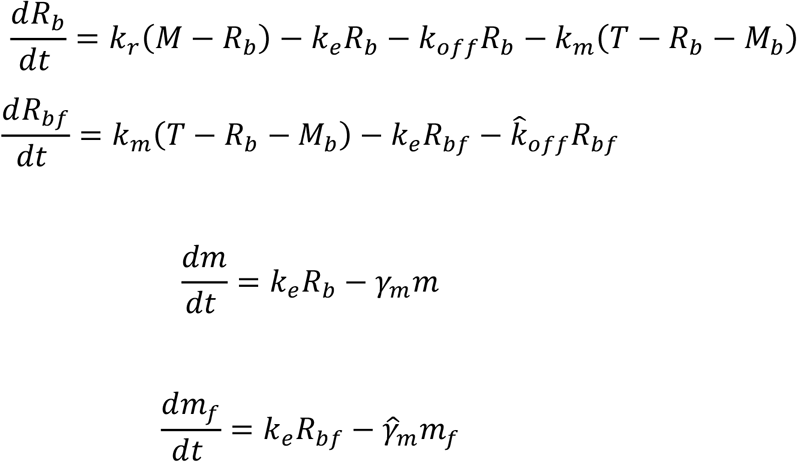

Here RNA polymerase II (*R*_*b*_) initiates transcription with rate *k*_*r*_ at *M* binding sites and elongates with rate *k*_*e*_ to form cytoplasmic mRNA (*m*), which subsequently decays with rate *γm*. Promoter-bound *R*_*b*_ can drop off at rate *k*_*off*_ and the CF is assumed to modulate this rate. The nuclear factor binds RNA polymerase with rate *k*_*m*_ to form a complex *R*_*bf*_ that elongates at the same rate, but at a different drop-off rate 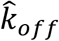. We assume that the factor imprints the nascent RNA complelex and is exported with it to the cytoplasm to create mRNA species *m*_*f*_. Once this factor-bound mRNA is degraded in the cytoplasm with rate 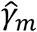, the factor is immediately imported back to the nucleus. *T* is the total intracellular level of the CF, which can be present in three forms: a free factor in the nucleus; bound to the elongating RNA polymerase complex; bound to mRNA in the cytoplasm. The transcription (synthesis rate) and mRNA stability impacts of the CF are modeled through parameters 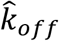 and 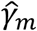. For example, an activator-only factor reduces the 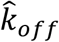 of the factor-bound polymerases, which now makes them more likely to complete transcription elongation and to export processes to synthesize cytoplasmic mRNA. An activator that destabilizes mRNA can be modeled by a reduced 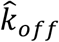 and an increased 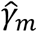 compared to *γ*_*m*_ and, similarly, a stabilizer reduces 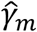.

To generate Figure 2A, we plot the net mRNA level (*m*_*f*_ + *m*) as a function of the initiation rate *k* _*r*_ for an activator that stabilizes mRNA 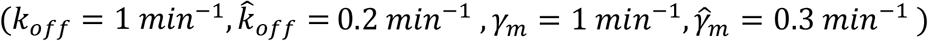, only the activator 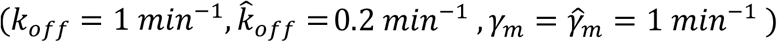, and an activator that destabilizes mRNA 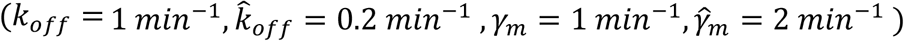. Other parameters can be taken, such as *T* = 3000, *M* = 1000, *k*_*e*_ = 1 *min*^*−*1^, *k*_*m*_ = 10 *min*^*−*1^.

In Figure 2B, we plot the net mRNA level as a function of time by solving the ODE model for turning an activator ON. We use the parameters in Figure 2A to consider synergistic regulation (activator that stabilizes mRNA), no crosstalk (only the activator) and buffering (the activator that destabilizes mRNA). In addition to the parameters in Figure 2A, we also consider *k* _*r*_ = 1 *min*^*−*1^ and the initial conditions of the ODE are taken as:

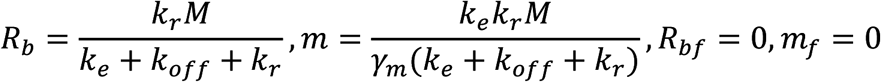

that correspond to the steady-state value of the ODE model in the absence of the activator. The net mRNA levels are normalized to their value at time *t* = 0.

In Figure 2C, we consider that a repressor is turned ON with four different cases: a repressor that destabilizes mRNA 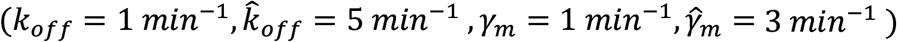, a repressor that only impacts transcription 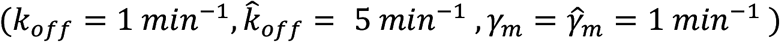, a repressor that only destabilizes mRNA 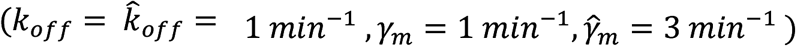 and a repressor that stabilizes mRNA 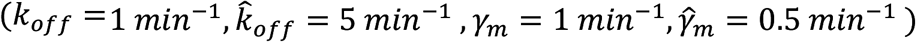. All the other parameters are as shown in Figures 2A-B, and the plotting procedure is as depicted in Figure 2B.

## Acknowledgements

We thank all the members of the GFL (Universitat de València) and Gene Expression (IBiS) Laboratories for their helpful discussion. This work was funded with grants PID2020-112853GB-C31, BFU2016-77728-C3-3-P and RED2018-102467-T to J.E.P-O., and BFU2016-77728-C3-1-P to S.C. all of them funded by MCIN/AEI/10.13039/501100011033 and grant BIO-271 from Junta de Andalucía to S.C.

## Data availability

The GRO data for the *sfp1* mutant are stored in the GEO repository (accession number GSE57467). The rest of the analyzed mutants datasets are from the ArrayExpress database (accession number E-MTAB-1525).

